# Carnitine deficiency alters fuel metabolism and voluntary wheel running in mice

**DOI:** 10.64898/2026.03.27.714765

**Authors:** Meagan S Kingren, Daniel G Sadler, Elijah Bolin, Izak Harville, James Sikes, Renny Lan, Elisabet Børsheim, Craig Porter

## Abstract

**Background:** Carnitine plays an obligatory role in energetics owing to its role in the translocation of long-chain fatty acids into the mitochondrion for oxidation. Here, we determined the metabolic and behavioral consequences of systemic carnitine deficiency (SCD) in mice.

**Methods:** Female C57BL/6J mice were randomized to receive normal drinking water (control, n = 8) or drinking water supplemented with mildronate 4g.L-1 (mildronate, n = 8) for 21 days. Body composition was assessed at baseline and post treatment. Metabolic and behavioral phenotyping was performed continuously over 72 hours following 14 days of control or mildronate treatment. Stable isotope were used to assess whole-body substrate oxidation. Carnitine subfractions were quantified in skeletal muscle and liver, as was mitochondrial respiratory function. Liver and muscle samples also underwent proteomic analysis.

**Results:** Mildronate treatment depleted total carnitine in muscle and liver by ∼97% (*P* < 0.001) and ∼90% (*P* < 0.001), respectively. Carnitine depletion was accompanied by lower total energy expenditure (*P* = 0.01), attributable to lower voluntary wheel running (*P* = 0.01). Oxidation rates of palmitate (*P* < 0.01) but not octanoate were lower whereas rates of glucose oxidation were greater in carnitine depleted mice (*P* < 0.01). Mitochondrial respiratory capacity was unaltered by carnitine deficiency. Carnitine deficiency remodeled muscle and liver proteomes to support lipid oxidation and energy production.

**Summary:** In mice, carnitine deficiency is characterized by decreased long-chain fatty acid oxidation despite preserved mitochondrial respiratory capacity. Carnitine deficiency resulted in lower voluntary exercise and a concomitant reduction in energy expenditure.

## Introduction

Carnitine, meaning *flesh* in Latin, was first identified in meat extracts at the turn of the 19^th^ century (1), and is now known to be a small water-soluble quaternary amine formed from the amino acids methionine and lysine (2). Around 95% of carnitine in the body is confined to striated muscle (3), where it plays two well-documented roles in bioenergetics (4, 5). First, as an essential co-factor for carnitine palmitoyl transferase I (CPT1), the rate limiting enzyme in long-chain fatty acid oxidation, carnitine is required for transport of long-chain fatty acids into mitochondria for oxidation (6, 7). Second, carnitine plays an important role as an acetyl group buffer (8, 9). Specifically, acetylation of free carnitine by the enzyme carnitine acetyl transferase (CAT) prevents acetyl-coA accumulation from depleting the mitochondrial co-enzyme A pool during increased flux through the pyruvate dehydrogenase complex (4).

Mitochondrial diseases are a common form of genetic disease, affecting approximately one in every 5,000 births (10). In particular, lipid storage myopathies resulting from the loss of a gene critical to the oxidative disposal of fatty acids are a prevalent manifestation of mitochondrial disease (11, 12). Loss of the organic cation transporter (OCTN2) responsible for maintaining plasma and tissue carnitine levels, which results in primary carnitine deficiency (PCD), is thought to be the most common myopathy resulting from a genetic (or primary) mitochondrial disease (10, 13–16). The loss of OCTN2, which is responsible for carnitine uptake from the intestines, retention of carnitine in renal tubules, and transport of carnitine from blood plasma into tissues, results in profound tissue carnitine depletion. The incidence of PCD ranges from 1:20,000 to 1:120,000 depending on geographical region, and results in a pronounced myopathy that includes excessive ectopic lipid storage in the liver and skeletal muscle, intolerance to physical activity, and rhabdomyolysis. In addition to PCD, secondary or acquired mitochondrial diseases can develop as part of other disease processes. Secondary carnitine deficiency (SCD) can develop in response to several chronic disease states including kidney disease requiring long-term dialysis (17–19), severe trauma (20), and chronic use of the anti-seizure medication valproate (21, 22). Although not associated with the loss of OCTN2, SCD results in tissue carnitine depletion, and incurs similar symptoms to those accompanying PCD. Accordingly, carnitine deficiency represents a mitochondrial disease that can manifest from a primary genetic origin (PCD), or one that can be acquired as the result of chronic illness (SCD).

Currently there are limited platforms to study PCD or SCD beyond observational studies in patients. The juvenile visceral steatotic (JVS) mouse has a spontaneous loss of the OCTN2 transporter (23), and is viable if supplemented with carnitine from weaning. However, the extreme phenotype of the JVS mouse has limited use in terms of a preclinical model of carnitine deficiency. The lack of strong preclinical models of carnitine deficiency means that there is a paucity of data concerning the metabolic impact of tissue carnitine deficiency. Further, the absence of efficacious treatment strategies for carnitine deficiency is symptomatic of the lack of robust preclinical models of PCD/SCD. We propose leveraging the commercially available drug mildronate (24), known to inhibit hepatic carnitine biosynthesis, renal carnitine reabsorption, and tissue uptake (25) of carnitine to develop, and comprehensively phenotype, a mouse model of carnitine deficiency. This model will provide a powerful tool to better understand the metabolic consequences of carnitine deficiency, as well as a platform to test the efficacy of interventions aimed at improving metabolism and function in patients with PCD/SCD.

## Materials and Methods

### Animals

The present study was approved by the Institutional Animal Care and Use Committee and the Institutional Biosafety Committee at the University of Arkansas for Medical Sciences. Sixteen female C57Bl/6J (#000664, Jackson Laboratories, Bar Harbor, ME, USA) mice (∼8 weeks old) were individually housed at 24°C on a reverse light cycle (light 7pm – 7am) with *ad libitum* access to a standard chow diet (TD.95092 [18.8% protein, 17.2% kcal fat, 63.9% kcal carbohydrate, 3.8 kcal/g], Envigo Teklad Diets, Madison, WI, USA). Mice were acclimated at our facility for ∼1 week prior to being randomized to either a control group (CON, n = 8) or mildronate group (MIL, n = 8). Mice in the CON group received purified drinking water (Milli-Q Advantage A10 system, Billerica, MA, USA) for 4 weeks, while mice in the MIL group received purified drinking water supplemented with mildronate (Shandong Bangda Pharmaceutical Co, ShanDong, China) at a concentration of 4 mg/mL to achieve a dose of ∼ 800 mg · kg^-1^ · day^-1^. Fresh mildronate solutions were prepared every 7 days.

### Body Composition Analysis

Body composition was measured by quantitative magnetic resonance imaging (qMRI) using EchoMRI-1100 (EchoMRI, Houston, Texas, USA). Fat-free mass (FFM) was calculated as the difference between body weight and fat mass (FM). Body composition was determined prior to randomization to CON and MIL (i.e., baseline), and then again ∼ 3-weeks later.

### Behavioral and Metabolic Phenotyping

After 14 days of randomization to CON or MIL, mice underwent metabolic and behavioral phenotyping. Mice were individually housed for 96 hours in specialized cages that permit rates of oxygen consumption (V̇O_2_) and carbon dioxide production (V̇CO_2_) to be measured (Sable Systems International, Las Vegas, NV, USA), from which energy expenditure (EE) was calculated. During this time, food and water intake, activity, and voluntary wheel running were also continuously recorded. EE was calculated per the Weir equation:

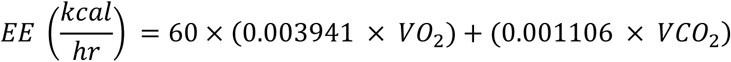

Data files from metabolic and behavioral phenotyping studies were processed using one-click-macros to distill data into hourly averages/totals and provided circadian reports for each 12-hour light cycle (Sable Systems International, Las Vegas, NV, USA). Total EE (TEE) was calculated by summing average rates of EE for each light cycle, where 24-TEE is the sum of TEE determined for both the light and dark cycle (kcal/cycle). Basal EE (BEE) was calculated from the 30-minute period with the lowest average EE (kcal/hour) in the light cycle and extrapolated to 24 hours. Resting EE (REE) was calculated from the 30-minute activity period with the lowest average EE (kcal/hour) in the light cycle and extrapolated to 24 hours. EE of individual wheel running bouts was calculated as the EE exceeding the BEE associated with each individual running bout. Total daily wheel EE was calculated by summing the EE of all wheel running bouts for each day and averaging across the measurement period. Peak activity EE (Active EE) was calculated as the highest EE (kcal/hour) over a 15-minute period in the dark cycle. All data were averaged over two consecutive 24-hour periods. The measurement of cage ambulation that included all gross and fine movements but excluded any meters ran directly on the running wheel is termed all meters.

Mice remained individually housed in metabolic phenotyping cages (Sable Systems International, Las Vegas, NV, USA) for a further 3 days following metabolic and behavioral phenotyping to undergo stable isotope studies to assess fatty acid and glucose oxidation. Briefly, we performed three separate stable isotope studies over three consecutive days, where animals received an intraperitoneal injection of either 1-^13^C glucose (99%), U-^13^C_16_ palmitate (99%) and 1-^13^C octanoic acid (99%) on separate days, to quantify the impact of carnitine depletion on CPT1 function and carbohydrate flux *in vivo*. At approximately 9 am each morning following a ∼ 2 hr. fast, animals received an I.P. injection of either 1-^13^C glucose, U-^13^C_16_ palmitate, or 1-^13^C octanoate (Cambridge Isotopes Laboratories Inc., Tewksbury, MA, USA). Isotopes were dissolved in sterile 0.09% saline (glucose (10 mg/mL) or octanoate (4 mg/mL)) or 30% bovine serum albumin (palmitate (1.5 mg/mL). For each isotope, a volume of 250 µL was injected I.P., representing doses of ∼ 125, 50 and 18.75 µg/g for glucose, octanoate, and palmitate, respectively (assuming a 20 g mouse). Animals were housed in the fasted state for up to 3 hours after isotope injections during which ^13^CO_2_ enrichment in the expired breath was continuously measured.

### Tissue Collection

Mice were euthanized by inhalation of a rising CO_2_ concentration at a displacement rate of 10% to 30% volume/min until all movement ceased, followed by an additional 1-2 min in the chamber. Euthanasia was confirmed by cervical dislocation and/or pneumothorax. The heart, liver, m. soleus, and m. gastrocnemius were collected. Portions of liver and m. gastrocnemius were placed in an ice-cold preservation (BIOPS) buffer (2.77 mM CaK _2_ EGTA, 7.23 mM K_2_ EGTA, 20 mM imidazole, 20 mM taurine, 50 mM MES hydrate, 0.5 mM dithiothreitol, 6.56 mM MgCl_2_ ·6H_2_ 0, 5.77 mM Na_2_ ATP, and 15 mM Phosphocreatine; pH 7.1) and transferred to the laboratory for high-resolution respirometry (HRR).

### Measurement of Tissue Metabolites

#### Triglyceride Concentration

The triglyceride concentration in the liver and heart tissue, respectively, was quantified spectrophotometrically using a commercially available kit (MAK266, Sigma-Aldrich). Tissue samples were homogenized in 5% Nonidet P-40 Substitute and subjected to two cycles of heating at 90°C for 3 min followed by cooling to room temperature. After centrifugation at maximum speed to remove insoluble material, supernatants were diluted 10-fold and incubated with lipase for 20 min at room temperature to hydrolyze triglycerides to glycerol and fatty acids. Samples were then incubated with the enzyme mix and probed at room temperature for 60 min, protected from light. Absorbance was measured at 570 nm using a microplate reader, and triglyceride concentrations were calculated from a standard curve.

#### Carnitine Moieties

All stock standards (1 mg/mL) were prepared in ethanol. L-Carnitine, Acetylcarnitine (C2), Propionylcarnitine (C3), Butyrylcarnitine (C4), Isovalerylcarnitine (iC5), Caproylcarnitine (C6), Octanoylcarnitine (C8), Decanoylcarnitine (C10), and Lauroylcarnitine (C12) were purchased from Sigma Aldrich (St. Louis, MO). Myristoylcarnitine (C14), Palmitoylcarnitine (C16), Stearoylcarnitine (C18), Oleoylcarnitine (C18:1), Linoleoylcarnitine (C18:2), D3-C14-Carnitine, D3-C16-Carnitine, and D3-C18:1-Carnitine were purchased from Cayman Chemical (Ann Arbor, MI). D9-L-Carnitine was purchased from Cambridge Isotope Laboratories (Tewksbury, MA), while D3-C3-Carnitine and D9-iC5-Carnitine was purchased from Sigma Aldrich (St. Louis, MO). All solvents were LCMS grade from Fisher Scientific (Pittsburgh, PA). Working calibration standards (0-20,000 ng/mL, 15 points) were diluted in 30% aqueous methanol and then spiked with isotope labelled internal standard mix (ISTD; 500 ng/mL).

Approximately 75 mg liver or muscle tissue was spiked with 100 µL of ISTD (500 ng/mL) before being homogenized in 1 mL of cold acetonitrile:methanol (3:1) using a Bertin Precellys24 homogenizer at 5000 rpm for 2 cycles of 20 seconds. Samples were cooled on dry ice for 2 min, and homogenization repeated. Homogenates (70 mg/mL) were mixed at 1000 rpm and 4°C for 20 min before centrifuging at 18,000 x g and 4°C for 10 min. Supernatant (1 mL) was dried down in a vacuum concentrator, then resuspended in 300 µL of 30% methanol using vortex for 20 seconds and sonication bath for 5 min. The reconstitution was centrifuged at 18,000 x g and 4°C for 5 min, then 50 µL of the supernatant was transferred to an LC vial for LC-MS analysis. 10 µL of each supernatant was also used to make a pooled sample for quality control (QC).

All mass spectrometry instrumentation was from Waters, Corporation (Milford, MA). Chromatographic separation was performed on an ACQUITY Premier UPLC fitted with an XSelect CSH C18 reverse-phase column (100 x 2.1 mm, 2.5 µm) kept at 30°C while samples were kept at 4°C. Five µL of each sample was injected and processed at the flow rate of 200 µL/min. Mobile phases consisted of 0.1% trifluoroacetic acid in water (A) and 0.1% trifluoroacetic acid in acetonitrile (B) with a 20 min elution gradient as follows: hold 30% B 1 min, ramp to 65% B over 3 min, hold 65% B 3 min, ramp to 80% B over 2 min, hold 80% B for 2 min, ramp to 95% B over 2 min, hold 95% B 4 min, return to 30% B in 1 min, hold 30% B until 20 min.

The quantification of free carnitine and acylcarnitines was carried out on a SELECT SERIES Cyclic IMS using time-of-flight (ToF) functionality in positive ionization mode. Data acquisition and quantitation was performed using MassLynx V4.2 and UNIFI 1.9.13 software, respectively. Prior to use, the Waters TOF G2-S sample kit was used for instrument calibration. MSe data was acquired in V-mode positive electrospray ionization at a mass range of 50-600 Da. Capillary, cone, and source offset were set to 2.5 kV, 30 V, and 10 V, respectively. Desolvation gas flow and temperature were at 800 L/hr and 550°C, respectively, while the source temperature was 120°C. Trap and transfer collision energies were set to 6 V and 4 V, respectively. Accurate ion masses were extracted from the Multiple Stage Exponentially-Sampled (MSE) data at low energy state (4 V) for quantification, while multiple fragment ions for each target were extracted at high energy state (6 V) for compound verification.

#### High-Resolution Respirometry (HRR)

Tissue samples were transferred to the laboratory immediately after collection for HRR. Approximately 5-10 mg of soleus muscle was permeabilized with 5μM saponin for 10-15 min at 4°C. HRR was carried out on approximately 2-4 mg of tissue in an Oroboros O2k respirometer chamber (Oroboros Instruments, Insburck, Austria) containing 2 mL buffer (0.5 mM EGTA, 3 mM MgCl_2_ ·6H_2_ 0, 60 mM lactobionic acid, 20 mM taurine, 10 mM KH_2_ PO_4_, 20 mM HEPES, 110 mM sucrose, and 1 g/L bovine serum albumin). Liver tissue was minced in BIOPS buffer before being blotted on filter paper briefly prior to being weighed. Approximately 1-2 mg (wet weight) of liver was transferred into the chamber of an Oxygraph-2K (O2k) high-resolution respirometer (Oroboros Instruments, Innsbruck, Austria) containing 2 ml of buffer (MiR05 composition: 0.5 mM EGTA; 3 mM MgCl2; 0.5 M K–lactobionate; 20 mM taurine; 10 mM KH_2_PO_4_; 20 mM HEPES; 110 mM sucrose; and 1 mg/mL essential fatty acid free bovine serum albumin) for assessment of mitochondrial bioenergetics. HRR analysis was performed on the same day of sample collection, typically within 2-4 hr. of euthanasia. Temperature was maintained at 37°C and O_2_ concentration within the range of 200–400 nmol/mL for all respirometry analyses. O_2_ concentration within the Oxygraph chamber was recorded at 2–4 s intervals (DatLab, Oroboros Instruments, Innsbruck, Austria) and used to calculate respiration per milligram of wet tissue weight.

All respiratory measures were made sequentially in the same sample. First, 50 μM palmitoyl-CoA and 50 mM malate were added to the chamber to record leak respiration supported primarily by the electron transferring protein. This was followed by the addition of 5 mM pyruvate, 2 mM malate, and 10 mM glutamate to record leak respiration supported primarily by Complex I. Thereafter, respiration was coupled to ATP production by the addition of ADP (7.5 mM). Coupled respiration was then assayed after the addition of the Complex II substrate succinate (10 mM). Next, 10 mM glycerol-3-phosphate was added to the chamber to assay coupled respiration supported by Complex I, Complex II, and glycerol-3-phoshate dehydrogenase. Respiration was then assayed in the uncoupled state following titration of carbonyl cyanide m-chlorophenyl hydrazone (CCCP) in 0.5 μM increments (up to a final concentration of 2.5 μM). Electron transfer was then inhibited by the addition of antimycin A (2.5 μM). The respiratory capacity of Complex IV was subsequently assayed following the addition of 800 mM ascorbate and 200 mM N,N,N’,N’-Tetramethyl-*p*-phenylenediamine dihydrochloride. Finally, Sodium Azide (300mM) was added the chamber to inhibit Complex IV. Flux control ratios were calculated by normalizing to maximal uncoupled respiration.

#### Quantitative Proteomics

Quantitative proteomics were carried out at the IDeA National Resource for Quantitative Proteomics as previously described (26, 27). Briefly, total protein from tissue (n = 8 per group per tissue) was concentration-matched, reduced, alkylated, and purified by chloroform/methanol extraction prior to digestion with sequencing grade modified porcine trypsin (Promega, Madison, WI, USA). Tryptic peptides were then separated by reverse phase XSelect CSH C18 2.5 um resin (Waters, Milford, MA) on an in-line 150 x 0.075 mm column using an UltiMate 3000 RSLCnano system (Thermo). Using a 60 min gradient from 98:2 to 65:35 buffer A:B ratio (Buffer A = 0.1% formic acid, 0.5% acetonitrile; Buffer B = 0.1% formic acid, 99.9% acetonitrile), peptides were eluted and subsequently ionized by electrospray (2.4kV) and mass spectrometric analysis on an Orbitrap Exploris 480 mass spectrometer (Thermo). To assemble a chromatogram library, six gas-phase fractions were acquired on the Orbitrap Exploris with 4 m/z Data-Independent Acquisition (DIA) spectra (4 m/z precursor isolation windows at 30,000 resolution, normalized Automatic Gain Control (AGC) target 100%, maximum inject time 66 ms) using a staggered window pattern from narrow mass ranges using optimized window placements. Precursor spectra were acquired after each DIA duty cycle, spanning the m/z range of the gas-phase fraction (i.e., 496-602 m/z, 60,000 resolution, normalized AGC target 100%, maximum injection time 50 ms). For wide-window acquisitions, the Orbitrap Exploris was configured to acquire a precursor scan (385-1015 m/z, 60,000 resolution, normalized AGC target 100%, maximum injection time 50 ms) followed by 50 x 12 m/z DIA spectra (12 m/z precursor isolation windows at 15,000 resolution, normalized AGC target 100%, maximum injection time 33 ms) using a staggered window pattern with optimized window placements. Precursor spectra were acquired after each DIA duty cycle.

#### Proteomic Data Analysis

Following acquisition, data were searched using an empirically corrected library, and a quantitative analysis was performed to obtain a comprehensive proteomic profile. Proteins were identified and quantified using EncyclopeDIA and visualized with Scaffold DIA using 1% false discovery thresholds at both the protein and peptide level (28). Protein exclusive intensity values were assessed for quality and normalized using ProteiNorm (29). The data was normalized using Cyclic Loess and statistical analysis was performed using Linear Models for Microarray Data (limma) with empirical Bayes (eBayes) smoothing to the standard errors (30). For pathway analysis, proteins with a *P*-value <0.05 were considered significant. Differentially abundant proteins (DAPs) were then converted from their UniProt ID (protein) to Ensembl ID (mouse gene) for pathway analysis. Using Ingenuity Pathway Analysis software, the Ensembl IDs of DAPs were subjected to canonical pathway analysis (28). Volcano plots were generated in R 4.2+ using EnhancedVolcano. UpSetR was used to generate the UpSet plot (29).

#### Statistical Analysis

Primary outcome measures for this study were EE and substrate oxidation data. Based on our previously published data (30), where we reported a ∼15% difference in whole body substrate oxidation rates (using indirect calorimetry) in carnitine depleted rats compared to normal rats, we estimated needing to study n = 7 animals per group to achieve a power of 0.8 and an alpha of 0.05 in order to detect true differences in group means. In total, we studied n = 8 animals per group. Animals were randomized to control or mildronate groups by body mass. To mitigate against any possible environment effects (e.g., temperature, humidity, light, and noise gradients), cage placement within a cage racks was randomized when possible. The order in which animals were euthanized was also randomized. All values are presented as means with standard errors. Comparisons between groups were tested using an unpaired t-test (Prism, version 10; GraphPad, San Diego, CA, USA).

## Results

### Animals

Characteristics of all animals are presented in **Table 1**. During the first 21 days of the study period, food and water were comparable (**Table 1**). Growth rate during the study period was also comparable between CON and MIL groups (**Table 1**). At the conclusion of the 4-week study, body masses were similar between CON and MIL groups (**Table 2**). Absolute liver and heart masses were also not different between CON and MIL groups; however, the relative mass of both the liver and heart were greater in animals in the MIL group versus CON (**Table 2**). The triglyceride concentration of the liver and heart were also greater in animals in the MIL group versus CON, although this only reached statistical significance in the liver (**Table 2**).

**Table 1.** Animal feeding, weight gain, and mildronate intake. Data from animal home cages from day 1 to 21 of randomization to Control (CON) or Mildronate (MIL), where mice were singly house but did not have access to wheels, are shown. Values are group means ± standard error, n = 8 per group.

|  | CON | MIL |
| --- | --- | --- |
| Body Mass Change (g/day) | $0.036 \pm 0.011$ | $0.037 \pm 0.015$ |
| Water Intake (mL) | $4.1 \pm 0.2$ | $3.8 \pm 0.3$ |
| Food Intake (g) | $3.5 \pm 0.2$ | $3.7 \pm 0.2$ |
| Mildronate Intake ($\text{mg} \cdot \text{kg}^{-1} \cdot \text{day}^{-1}$) | 0 | $790 \pm 41$ |

**Table 2.** Necropsy and tissue triglyceride data for mice randomized to Control (CON) or Mildronate (MIL). Values are group means ± standard error, n = 8 per group.

|  | CON | MIL | *P* |
| --- | --- | --- | --- |
| Body Mass (g) | $20.0 \pm 0.5$ | $19.3 \pm 0.6$ | $P > 0.05$ |
| Liver Mass (mg) | $902 \pm 66$ | $1042 \pm 47$ | $P > 0.05$ |
| Relative Liver Mass (%) | $4.5 \pm 0.3$ | $5.4 \pm 0.2$ | $P = 0.02$ |
| Liver Triglyceride (ng/uL) | $42.6 \pm 13.1$ | $103.0 \pm 17.2$ | $P = 0.02$ |
| Heart Mass (mg) | $121.1 \pm 5.4$ | $141.9 \pm 9.3$ | $P = 0.09$ |
| Relative Heart Mass (%) | $0.61 \pm 0.02$ | $0.74 \pm 0.04$ | $P = 0.03$ |
| Heart Triglycerides (ng/uL) | $11.2 \pm 3.5$ | $44.1 \pm 16.0$ | $P = 0.08$ |

### Metabolic and Behavioral Phenotyping

Continuous EE, wheel running behavior, and feeding behaviors for CON and MIL groups are reported over a 72-hour period and presented in **Figure 1**. Daily averages for TEE and its components, as well as movement parameters (ambulation and wheel running), are shown in **Figure 2**, where the averages of values from three consecutive days are presented.

**Figure 1.**
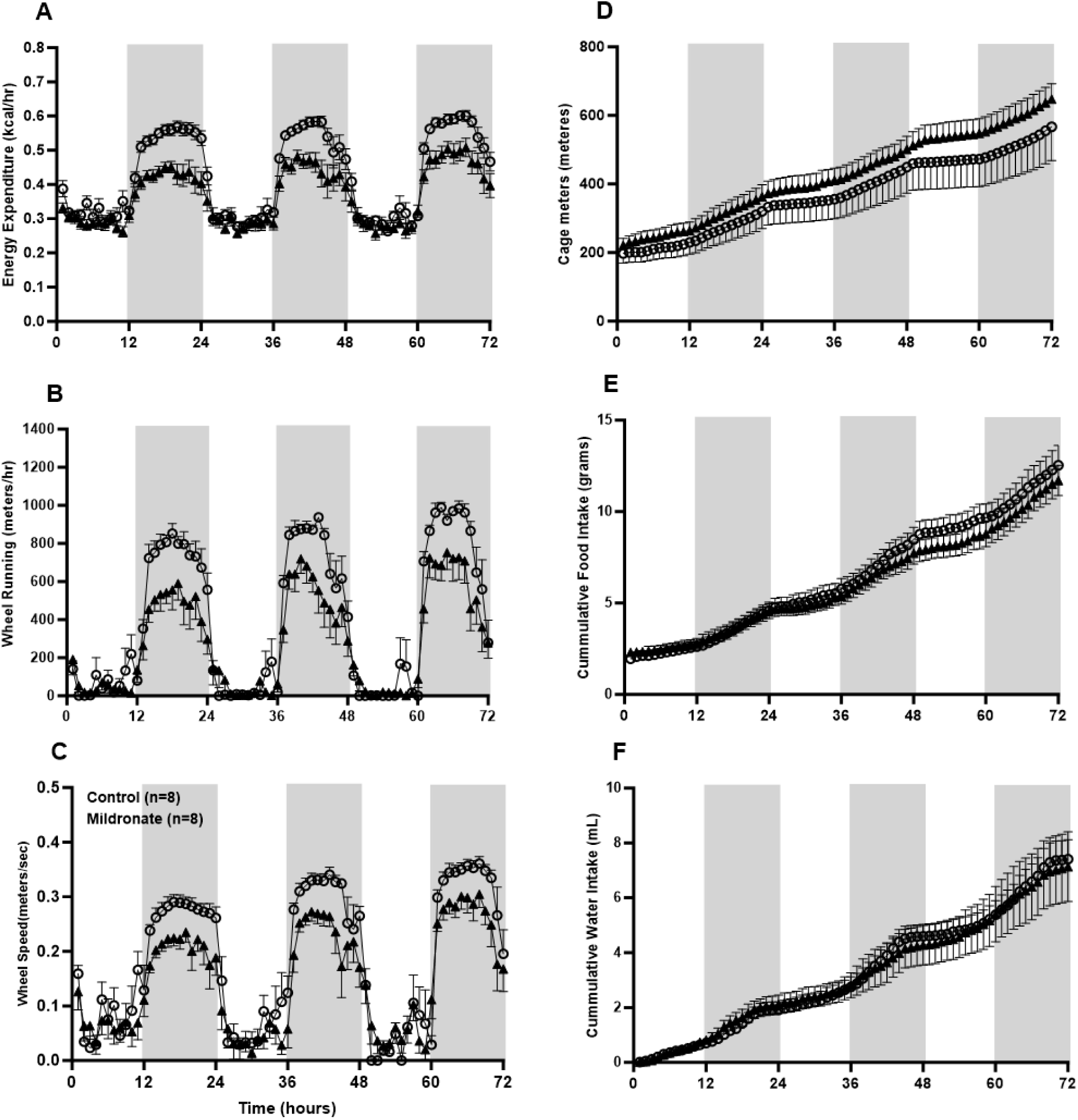
72 hours of continuous metabolic and behavioral phenotyping performed after two weeks of randomization to either Control (CON, open symbols) or Mildronate (MIL, closed symbols). Panel A shows energy expenditure data over time in CON and MIL treated mice. Voluntary wheel running distance (m/hr) and speed (m/s) in CON or MIL-treated mice are shown in Panels B and C, respectively. Panel D shows cumulative cage ambulation in meters, while Panels E and F show cumulative food and water intake, respectively. Gray shading represents the 12-hr dark period, where lights in the cabinets housing the mice were not illuminated. Values are group means ± standard error, n = 8 per group.

**Figure 2.**
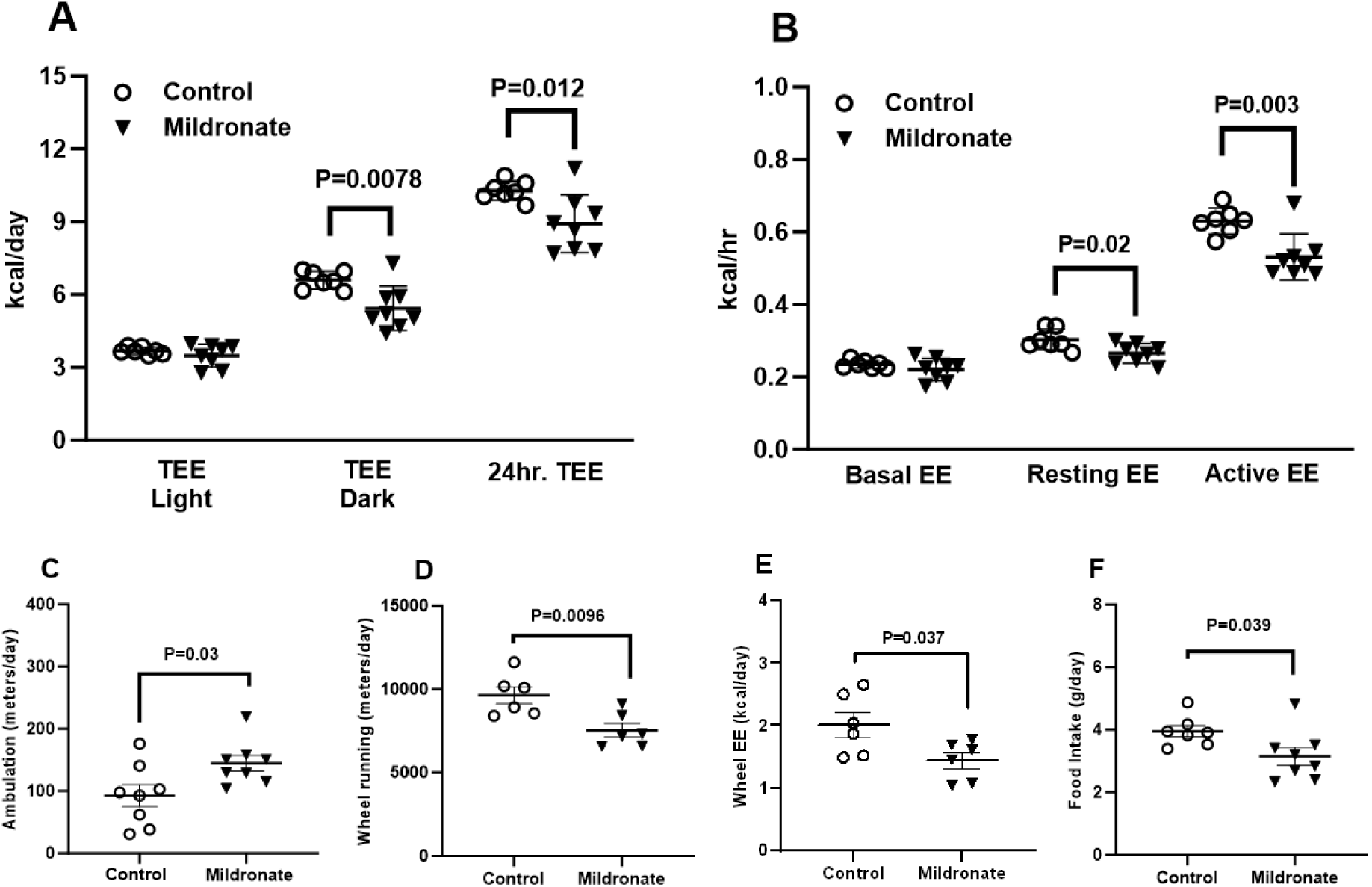
Metabolic and behavioral phenotyping outcomes performed after two weeks randomization to either Control (CON, open symbols) or Mildronate (MIL, closed symbols). Panel A shows total energy expenditure (TEE) of CON and MIL-treated mice in the 12-hour light and dark photo periods. 24-hour TEE is the sum of TEE in both the light and dark photo periods. Panel B shows energy expenditure in the basal, resting, and active states. Basal EE (BEE) was calculated from the 30-min period with the lowest average EE (kcal/hour) in the light cycle and extrapolated to 24 hours. Resting EE (REE) was calculated from the 30-min activity period with the lowest average EE (kcal/hour) in the light cycle and extrapolated to 24 hours. Peak activity EE (Active EE) was calculated as the highest EE (kcal/hour) over a 15-min period in the dark cycle. Panel C show daily cage ambulation (excluding meters ran on wheels). Panel D shows voluntary wheel running distance and Panel E shows energy expenditure related to wheel running. Total daily wheel EE was calculated by summing the EE of all wheel running bouts for each day and averaging across the measurement period. Panel F shows daily food intake. Values are group means ± standard error, n=8 per group.

There was an apparent separation in hourly EE rates in the dark period only (**Figure 1A**), where EE was greater in CON animals versus MIL. TEE, calculated by summing average rates of EE for each light cycle, was significantly greater in CON versus MIL in the dark period (*P* = 0.0078, **Figure 2A**), but not the light period. 24-hr TEE, representing the sum of TEE in both the light and dark cycles (kcal/day), was also significantly greater in CON versus MIL (*P* = 0.012, **Figure 2A**). BEE representing the lowest average EE (kcal/hour) over a 30 min period in the light cycle was not different between CON and MIL (**Figure 2B**). Resting EE (REE), representing the lowest average EE (kcal/hour) over a 30 min period with activity in the light cycle was significantly greater in CON versus MIL (*P* = 0.02, **Figure 2B**). Peak activity EE (Active EE) representing the highest EE (kcal/hour) over a 15-minute period in the dark cycle was also significantly greater in CON versus MIL (*P* = 0.003, **Figure 2B**).

In line with a separation in hourly EE rates in the dark period only, both wheel running volume (**Figure 1B**) and speed (**Figure 1C**) exhibited a similar pattern, where mice in the CON group ran more and ran faster than mice in the MIL group. Indeed, daily wheel meters was approximately 30% greater in CON mice compared to MIL mice (*P* = 0.0096, **Figure 2D**). Similarly, EE related to wheel running was also 30% greater in CON mice compared to MIL mice (*P* = 0.037, **Figure 2E**). Accordingly, greater TEE in CON versus MIL mice was largely attributable to greater EE caused by more wheel running (**Figure 2E**). While running volume was significantly greater in CON versus MIL mice (**Figure 1B and 2D**), cage ambulation was greater in the MIL group versus the CON group (*P* = 0.03, **Figure 1D and 2C**), presumably reflecting less time spent wheel running by mice in the MIL group. However, it is worth noting that this difference in ambulation represents ∼ 50 m/day (i.e. ∼ 150 versus 200 m/day, **Figure 2C**). So, while mice in the MIL group moved ∼ 50 m/day more than mice in the CON group in terms of ambulation around their cage, mice in the CON group moved 2000 m/day more than those in the MIL group overall due to greater voluntary wheel running (**Figure 2D**).

When housed in cages without access to running wheels, food intake in CON and MIL groups where similar (**Table 1**). Of interest, during metabolic and behavioral phenotyping experiments, where mice had access to running wheels, a modest separation in food intake was observed, with mice in the CON group consuming more food than those in the MIL group (**Figure 2E**). This modest increase in food intake was not accompanied by a measurable separation in water intake (**Figure 1F**). Greater food intake, amounting to ∼0.8 g/day greater food intake in CON versus MIL animals (**Figure 2F**), or around 3 kcal/day greater energy intake based on the energy density of the food animals were provided, likely reflects the greater wheel running and thus greater TEE of CON mice compared to MIL mice.

### Substrate Oxidation Rates

The oxidation rates of the long-chain fatty acid palmitate, glucose, and the medium-chain fatty acid octanoic acid are shown in **Figure 3**. The enrichment of ^13^CO_2_ in expired gas was greater in CON versus MIL mice following an i.p. injection of U-^13^C_16_ palmitate (P<0.01, **Figure 3A**), where the AUC for breath ^13^CO_2_ enrichment in the 2 hours following the administration of U-^13^C_16_ palmitate was significantly higher in CON compared to MIL (*P* = 0.014, **Figure 3B**). In contrast, the enrichment of ^13^CO_2_ in expired gas was greater in MIL versus CON mice following an i.p. injection of 1-^13^C glucose (*P* < 0.01, **Figure 3C**), where the AUC for breath ^13^CO_2_ enrichment in the 2 hours following the administration of 1-^13^C glucose was significantly higher in MIL compared to CON (*P* = 0.023, **Figure 3D**). There was no difference in the enrichment of ^13^CO_2_ in expired gas between CON and MIL mice following an i.p. injection of 1-^13^C octanoic acid (**Figure 3E and 3F**).

**Figure 3.**
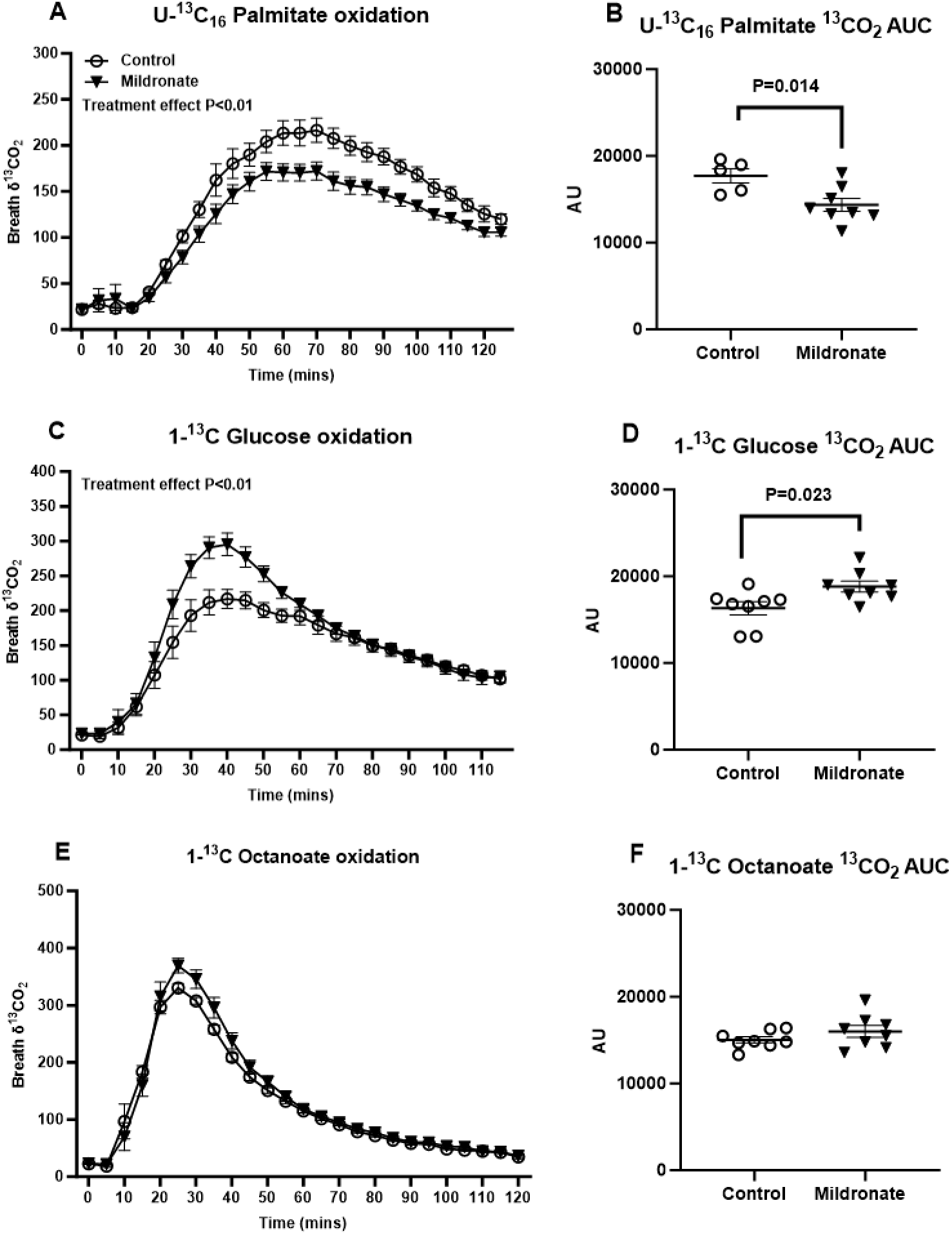
*In vivo* substrate oxidation rates following 3 weeks of randomization to either CON (open symbols) or MIL (closed symbols). Panel A shows breath ^13^CO_2_ enrichment over time following an i.p. injection of U-^13^C_16_ palmitate. The Area Under the Cure (AUC) for breath ^13^CO_2_ as a marker of palmitate oxidation is shown in Panel B. Panel C shows breath ^13^CO_2_ enrichment over time following an i.p. injection of 1-^13^C glucose. The AUC for breath ^13^CO_2_ as a marker of glucose oxidation is shown in Panel D. Panel E shows breath ^13^CO_2_ enrichment over time following an i.p. injection of 1-^13^C octanoate. The AUC for breath ^13^CO_2_ as a marker of octanoate oxidation is shown in Panel F. Values are group means ± standard error, n = 8 per group.

### Body Composition

Body mass and composition data at baseline and following approximately 3 weeks of either CON or MIL treatment are shown in **Figure 4**. Body mass significantly increased in both the CON (**Figure 4A**) and MIL (**Figure 4D**) groups during the study period, although this only reached statistical significance in the MIL group. Absolute fat mass declined significantly over the study in mice in the CON group (**Figure 4B**), but not the MIL group (**Figure 4F**). Similarly, relative fat mass declined significantly over the study in mice in the CON group (*P* = 0.016, **Figure 4C**), but not the MIL group (**Figure 4G**).

**Figure 4.**
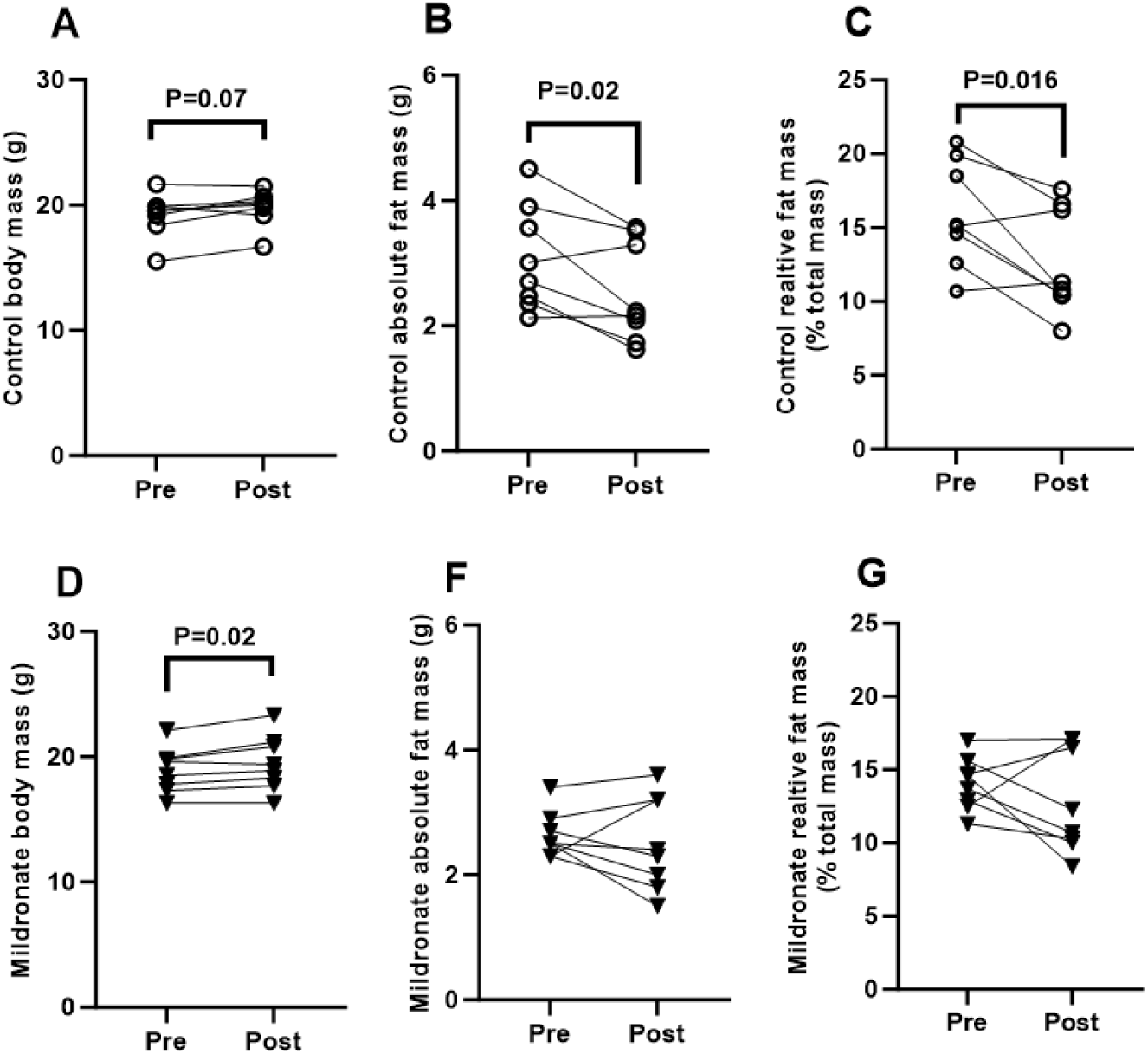
Body mass and composition data at baseline and following approximately 3 weeks randomization to either CON (open symbols) or MIL (closed symbols). Pre- and post- body mass, absolute fat mass and relative fat mass in CON mice are shown in Panels A, B, and C, respectively. Pre- and post- body mass, absolute fat mass and relative fat mass in MIL mice are shown in Panels D, E, and F, respectively. Pre and post values were compared within CON and MIL groups using a paired t-test. Values are group means ± standard error, n = 8 per group.

### Tissue Carnitine Concentrations

Concentrations of carnitine moieties in liver and gastrocnemius are presented in **Table 3** and **Table 4**, respectively. In the liver of MIL treated animals, free carnitine was 95% lower compared to CON (P<0.001). Acetyl-l-carnitine was not different in the liver of CON versus MIL animals. The concentrations of most long-chain acyl-carnitines (C14 to C18) were significantly lower in MIL versus CON treated mice, where the sum of long-chain acyl-carnitines moieties was ∼72% lower in MIL versus control (P<0.001). The concentration of the medium chain C6, C8 and C10 acyl-carnitines were significantly lower in the liver of MIL-treated mice compared to control; however, the concentration of C12 acyl-carnitine was not different between CON and MIL groups. The concentration of the short chain C5 and C3 acyl-carnitines were significantly lower in the liver of MIL-treated mice compared to control; however, the concentration of C4 acyl-carnitine was not different between CON and MIL groups. Total carnitine concentration, the sum of all carnitine moieties, was 90% lower in the liver of MIL treated mice versus CON mice.

**Table 3.** Liver Carnitine Concentrations. Total carnitine is the sum of free, acetyl and all acyl carnitine moieties. Values are group means ± standard error, n = 8 per group.

|  | CON | MIL | P |
| --- | --- | --- | --- |
| L-Carnitine (ng/mg) | 37.01 ± 16.77 | 1.66 ± 0.41 | < 0.001 |
| Acetyl-L-Carnitine (ng/mg) | 1.62 ± 0.52 | 1.94 ± 0.29 | 0.178 |
| C18-Carnitine (ng/mg) | 0.17 ± 0.15 | 0.07 ± 0.03 | 0.082 |
| C18:1-Carnitine (ng/mg) | 1.16 ± 0.43 | 0.25 ± 0.23 | < 0.001 |
| C18:2-Carnitine (ng/mg) | 0.57 ± 0.20 | 0.10 ± 0.07 | < 0.001 |
| C16-Carnitine (ng/mg) | 0.28 ± 0.09 | 0.06 ± 0.03 | < 0.001 |
| C14-Carnitine (ng/mg) | 0.02 ± 0.00 | 0.01 ± 0.01 | 0.208 |
| C12-Carnitine (ng/mg) | 8.78 ± 0.86 | 17.06 ± 4.61 | 0.12 |
| C10-Carnitine (pg/mg) | 11.19 ± 1.04 | 3.99 ± 1.49 | 0.002 |
| C8-Carnitine (pg/mg) | 8.09 ± 1.93 | 0.78 ± 0.73 | 0.005 |
| C6-Carnitine (pg/mg) | 7.40 ± 2.26 | 0.10 ± 0.09 | 0.009 |
| C5-Carnitine (pg/mg) | 18.88 ± 2.44 | 0.78 ± 0.41 | < 0.001 |
| C4-Carnitine (pg/mg) | 36.46 ± 3.87 | 29.57 ± 2.39 | 0.178 |
| C3-Carnitine (pg/mg) | 61.41 ± 9.77 | 8.32 ± 1.35 | < 0.001 |
| Total Carnitine (ng/mg) | 40.98 ± 17.28 | 4.15 ± 0.93 | < 0.001 |

**Table 4.** Skeletal Muscle (*m. gastrocnemius*) Carnitine Concentrations. Total carnitine is the sum of free, acetyl and all acyl carnitine moieties. Values are group means ± standard error, n = 8 per group.

|  | CON | MIL | P |
| --- | --- | --- | --- |
| L-Carnitine (ng/mg) | 40.95 ± 8.55 | 0.79 ± 0.23 | < 0.001 |
| Acetyl-L-Carnitine (ng/mg) | 13.65 ± 4.00 | 0.03 ± 0.01 | < 0.001 |
| C18-Carnitine (ng/mg) | 1.63 ± 1.21 | 0.14 ± 0.03 | < 0.01 |
| C18:1-Carnitine (ng/mg) | 4.45 ± 2.35 | 0.44 ± 0.05 | < 0.001 |
| C18:2-Carnitine (ng/mg) | 3.31 ± 1.81 | 0.20 ± 0.05 | < 0.001 |
| C16-Carnitine (ng/mg) | 1.48 ± 0.75 | 0.11 ± 0.01 | < 0.001 |
| C14-Carnitine (ng/mg) | 0.24 ± 0.13 | 0.03 ± 0.01 | < 0.001 |
| C12-Carnitine (pg/mg) | 78.22 ± 13.52 | 21.00 ± 1.63 | < 0.01 |
| C10-Carnitine (pg/mg) | 65.55 ± 9.37 | 21.40 ± 1.68 | < 0.001 |
| C8-Carnitine (pg/mg) | 51.26 ± 9.04 | 1.11 ± 0.20 | < 0.001 |
| C6-Carnitine (pg/mg) | 55.05 ± 7.62 | 8.95 ± 0.74 | < 0.001 |
| C5-Carnitine (pg/mg) | 59.06 ± 7.14 | 14.51 ± 1.11 | < 0.001 |
| C4-Carnitine (ng/mg) | 1.99 ± 0.34 | 0.02 ± 0.00 | < 0.001 |
| C3-Carnitine (pg/mg) | 78.62 ± 4.39 | 18.13 ± 1.44 | < 0.001 |
| Total Carnitine (ng/mg) | 68.08 ± 12.67 | 1.85 ± 0.18 | < 0.001 |

In the gastrocnemius of MIL-treated animals, there was a near complete depletion (98%) of free carnitine concentration compared to CON (*P* < 0.001). There was also a near complete depletion of acetyl-l-carnitine concentration (∼99%) in the skeletal muscle of the MIL group compared to control (*P* < 0.001). The concentrations of all long-chain acyl-carnitines moieties measured were markedly (∼ 90%) lower in MIL versus CON treated mice. Similarly, all medium chain (∼75%) short chain (∼ 97%) acyl-carnitines moieties were significantly lower in skeletal muscle of MIL treated mice compared to CON. Skeletal muscle total carnitine concentration was 97% lower in MIL treated mice versus CON Mice.

### High Resolution Respirometry

Mitochondrial respiration function data for the soleus muscle and liver are shown in **Figure 5**. There were no significant differences between CON and MIL groups for respiratory function in either the soleus (**Figure 5A**) or liver (**Figure 5B**). The respiratory control ratio for ADP and the substrate control ratios for both succinate and glycerol-3-phosphate were also similar in both the soleus and liver of CON and MIL groups (data not shown). In the soleus muscle, the flux control ratio for state 3 respiration supported by Complex I (0.52 ± 0.04 vs. 0.65 ± 0.02, *P* = 0.02), state 3 respiration supported by Complex I and Complex II (0.75 ± 0.04 vs. 0.86 ± 0.02, *P* = 0.03), and state 3 respiration supported by Complex I, Complex II, and glycerol-3-phosphate (0.081 ± 0.04 vs. 0.95±0.01, *P* = 0.007), were all significantly greater in the MIL group compared to CON (**Figure 5C**). In the liver, mitochondrial flux control ratios were similar between CON and MIL groups (**Figure 5D**).

**Figure 5.**
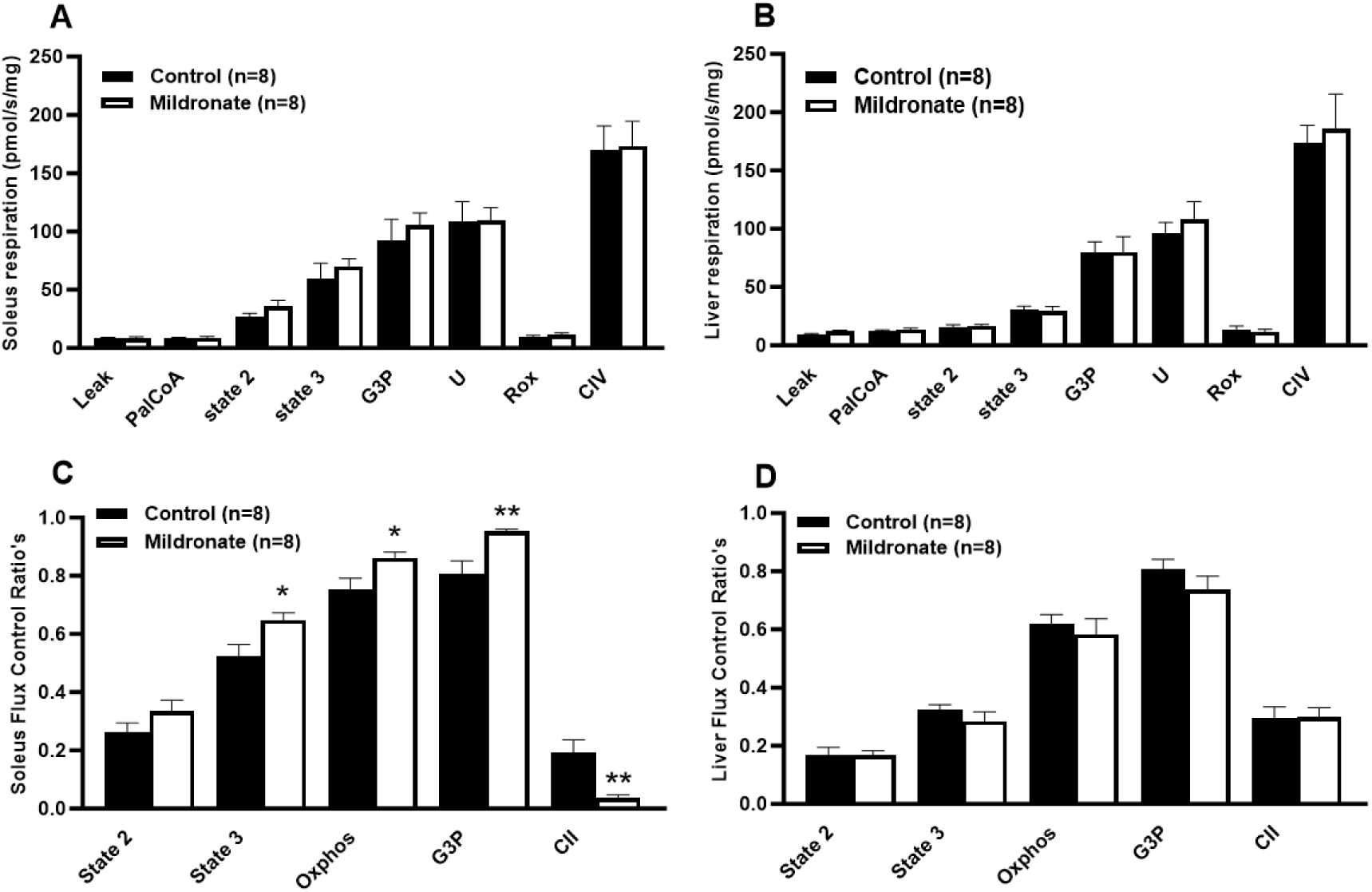
Mitochondrial respiration in permeabilized muscle and liver tissue following 4 weeks of randomization to either CON (black bars) or MIL (open bars). Panels A and B show respiratory flux per mg of tissue (wet weight) in the soleus muscle and liver, respectively. Panels C and D show respiratory flux control ratios for the soleus muscle and liver, respectively. Flux control ratios were calculated by normalizing specific respiratory rates to uncoupled (U) respiration. Values are group means ± standard error, n = 8 per group.

### Untargeted Proteomics

Untargeted proteomics data for the heart, liver, and soleus muscle are shown in **Figure 6**. In total, 699, 557, and 208 proteins were differentially abundant (DAPs) in the heart, liver, and soleus muscle from mice in the MIL versus CON groups, respectively (**Figure 6A**). Of these DAPs, 389 and 310 were up- and down-regulated, respectively, in the heart. In the liver, 245 and 312 DAPs were up- and down-regulated, while the soleus had 140 up- and 68 down-regulated DAPs, respectively. Ingenuity Canonical Pathway analysis of DAPs revealed that the MIL group demonstrated increases in mitochondrial fatty acid β-oxidation, regardless of tissue (**Figure 6B**). Indeed, 25 DAPs were similar across all three tissues, several of which related to fatty acid β-oxidation (**Figure 6C**). Both striated muscle, heart, and soleus muscle from the MIL group also demonstrated similar changes. Compared to the CON group, MIL heart and soleus demonstrated increases in the respiratory electron transport, oxidative phosphorylation, mitochondrial protein import, and Complex I biogenesis canonical pathways, as well as a decrease in the mitochondrial dysfunction pathway. In the liver, MIL supplementation resulted in increases in proteins related to lipid metabolism.

**Figure 6.**
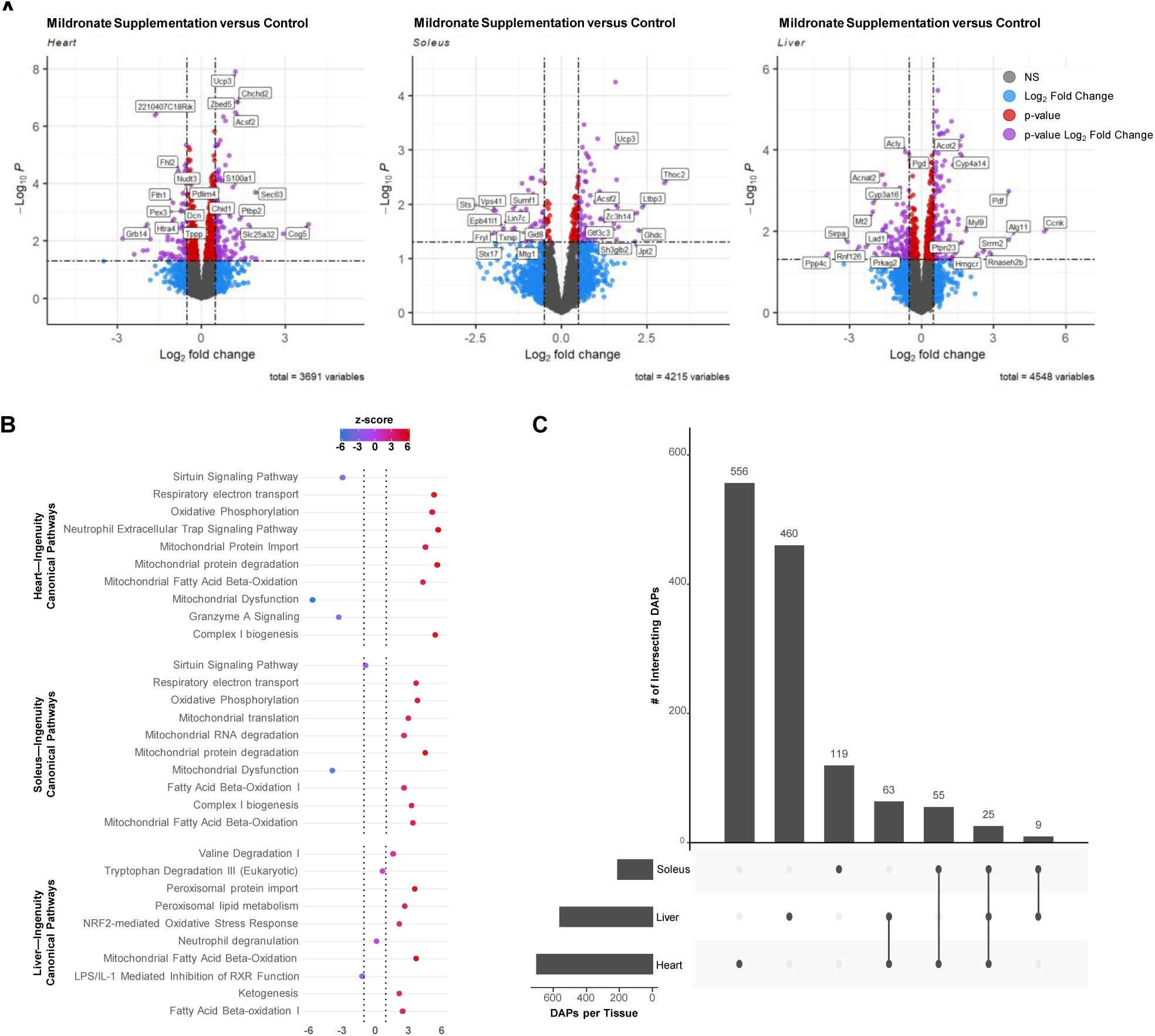
Untargeted proteomics in the heart, soleus, and liver mice randomized to Control (CON) or Mildronate (MIL). Panel A shows the volcano plots for each tissue, labeled with the 10 most up- and down-regulated DAPs. The top 10 most significantly altered Ingenuity Canonical Pathways for each tissue are shown in Panel B. The UpSet plot in Panel C shows the number of unique and overlapping DAPs for each tissue and tissue combination for MIL versus CON groups. Values are group means, n = 8 per group.

## Discussion

We set out to comprehensively phenotype a mouse model of carnitine deficiency to develop a platform to test the efficacy of interventions aimed at improving metabolism and function in patients with PCD/SCD. By combining physiological and behavioral phenotyping with stable isotope approaches, we show that tissue carnitine deficiency impairs long-chain fatty acid oxidation capacity despite preserved *ex vivo* mitochondrial respiratory function. Moreover, carnitine deficiency is associated with reduced voluntary exercise and a corresponding decline in energy expenditure, suggesting that systemic carnitine depletion alters muscle bioenergetics *in vivo*, with downstream effects on behavior and whole-body metabolism.

The butyrobetaine analogue mildronate is known to impair hepatic carnitine synthesis (25, 31), accelerate urinary carnitine excretion *in vivo* (25, 32), and reduce carnitine transport *in vitro* (33, 34). Previous rodent work has shown that mildronate reduces carnitine levels in the blood, heart, liver, and skeletal muscle (25, 30, 35–39). Here, we also show that 4 weeks of mildronate treatment markedly reduces all carnitine sub-fractions in skeletal muscle and liver, with total carnitine concentrations 90% lower in the liver and 97% lower in skeletal muscle compared to control mice. In particular, free carnitine – the predominant form of carnitine in resting muscle – was nearly completely depleted, with muscle free carnitine levels in mildronate-treated mice reduced to ∼2% of those in control mice. This severe depletion of free carnitine likely impaired flux through key carnitine-dependent enzymes, such as CPT1 and CAT, which rely on free carnitine as an essential acyl group carrier. In agreement with this, acetyl-l-carnitine and all long-chain acyl-carnitine moieties measured in the current study were markedly depleted in mildronate treated mice.

Despite marked carnitine depletion in skeletal muscle and liver of mildronate-treated mice, mitochondrial respiratory capacity remained intact in both tissues. Further, the Ingenuity Canonical Pathway for mitochondrial fatty acid β-oxidation was significantly upregulated in all three tissues for the MIL group. This suggests that the shift in fuel selection associated with carnitine deficiency does not arise from defects in the mitochondrial respiratory chain, but rather reflects impaired fatty acid transport into mitochondria upstream of β-oxidation. Interestingly, others have shown that 3-weeks of mildronate treatment does lower Complex II-supported state 3 respiration as well as maximal Complex IV respiratory flux in the rat soleus (40), a finding which we could not corroborate. However, it should be noted that this previous work assayed succinate-supported state 3 respiration in the presence of the Complex I inhibitor rotenone. It is possible that in the context of Complex I inhibition, Complex II-driven state 3 respiration is impacted by chronic carnitine deficiency. Nevertheless, this previous report by Bouitbir and colleagues (40) reported comparable state 3 respiration supported by Complex I in both the soleus and gastrocnemius of carnitine-deficient rats, in line with our observation in carnitine-deficient mice. While respiratory fluxes measured *ex vivo* appear largely unaffected in skeletal muscle of carnitine-deficient mice, we did observe increased flux control in the context of carnitine deficiency. This observation may reflect an adaptation in skeletal muscle that supports greater coupling of electron transfer to phosphorylation in the context of reduced entry of long-chain fatty acids into the mitochondrion. This is supported by increased canonical pathway expression for proteins related to respiratory electron transport, mitochondrial complex I biogenesis, and oxidative phosphorylation in the heart and soleus of mice in the MIL group versus CON.

Altering muscle carnitine availability (either increased or decreased carnitine availability) shifts skeletal muscle fuel selection and impacts muscle contractile function (19, 30, 38, 41–43). CPT1 is considered the rate limiting step in long-chain fatty acid oxidation. Mildronate-mediated carnitine deficiency has previously been shown to reduce the oxidation of radio-labelled palmitate in carnitine deficient rats (25), while indirect calorimetry has been used to show that in the fasted state, fat oxidation is attenuated while carbohydrate oxidation is accelerated in carnitine deficient rats (30). In this study, we used stable isotopes of palmitate and octanoic acid to assess how carnitine deficiency affects the oxidation of long- and medium-chain fatty acids, noting that entry of long-chain fats into mitochondria requires CPT1. We found that palmitate oxidation was significantly reduced in carnitine-deficient mice compared to controls, consistent with impaired mitochondrial import of long-chain fatty acids via CPT1. In contrast, octanoic acid oxidation was unaffected, aligning with the general view that medium-chain fatty acid oxidation is carnitine-independent. However, emerging data suggest this independence may be tissue-specific, with skeletal muscle and heart retaining some reliance on carnitine for medium-chain fatty acid oxidation. (44), which may explain the absence of any compensatory increase in octanoic acid in carnitine deficient mice. Our findings align with previous studies reporting reduced long-chain acylcarnitine accumulation and fatty acid oxidation under conditions of impaired CPT1 activity (45, 46), with *ex vivo* evidence showing muscle homogenates from systemic carnitine deficiency patients have a reduced capacity to oxidize long-chain fatty acids such as oleate and palmitate (13). In addition to impaired long-chain fatty acid oxidation, we observed a concurrent increase in the oxidation of isotopically labelled glucose in carnitine-deficient mice. This is in line with evidence of carnitine deficiency accelerating muscle carbohydrate use, and underscores the impact of severe carnitine deficiency on long-chain fatty acid oxidation and subsequently the integration of lipid and carbohydrate metabolism.

There is limited data regarding the impairment of skeletal muscle function in patients with documented muscle carnitine deficiency, such as those with end stage renal failure (17, 18, 47) or systemic carnitine deficiency patients (13, 14, 48). However, muscle carnitine deficiency is associated with exercise intolerance and reduced cardiorespiratory exercise capacity. In carnitine deficient rats, *ex vivo* peak isometric tension in the fast twitch extensor digitorum longus muscle is diminished in carnitine deficient rats (38). Further, during a treadmill running performance test, distance to exhaustion was 39% lower in carnitine deficient rats versus control (39), supporting the notion that altered fuel selection in the context of carnitine deficiency perturbs muscle energetics and ultimately function *in vivo*. In agreement with this, we observed a marked reduction in voluntary wheel running (∼30%) in carnitine-deficient mice. This was accompanied by a significant reduction in TEE. These findings underscore the physiological impact of carnitine deficiency, highlighting its ability to reduce volitional activity and total energy expenditure—suggesting that compromised muscle energetics in this context influences whole-body behavior and metabolism. Mice were only housed in cages with access to running wheels for the last 7 days of this 4-week study. While lower TEE attributable to running volume in carnitine-deficient animals was accompanied with lower food intake relative to control mice, lower TEE (as well as marginally albeit significantly lower resting EE) in carnitine-deficient mice may explain why adiposity did not change in carnitine-deficient mice while adiposity (both absolute and relative fat mass) declined in control mice during the 4-week study. Accordingly, prolonged carnitine deficiency and subsequent alterations in whole-body energetics may impact adiposity *in vivo*. This is broadly in agreement with human data indicating that the augmenting muscle carnitine content increases energy expenditure (41).

We employed an untargeted proteomic approach to determine whether changes in fuel oxidation and exercise behaviors brought about by carnitine deficiency were associated with compensatory adaptations in the proteome of muscle and liver. Overall, carnitine deficiency resulted in significant remodeling of the heart, liver, and skeletal muscle proteomes, with canonical pathway analysis indicating increases in fatty acid metabolism and oxidation, presumably to counter deficits in mitochondrial fatty acid import owing to carnitine deficiency. Of interest, in addition to enrichment of pathways involved in fatty acid oxidation, there was also enrichment in pathways involving branched chain amino acid oxidation and oxidative phosphorylation in heart and skeletal muscle of carnitine-deficient mice. The liver proteome demonstrated upregulation of pathways related to ketogenesis and peroxisomal lipid metabolism, along with elevated fatty acid β-oxidation, together indicating a possible compensatory response of the liver during carnitine deficiency. Indeed, in line with our mitochondrial flux control data, the remodeling of the muscle proteome is indicative of an adaptive response to support increased fuel catabolism in the context of diminished capacity for long-chain fatty acid trafficking.

In summary, we report that mildronate-mediated carnitine deficiency in mice is characterized by decreased capacity for long-chain fatty acid oxidation despite preserved mitochondrial respiratory capacity and an apparent compensatory increase in pathways involved in lipid metabolism, At the whole-body level, carnitine deficiency and reduced capacity for fatty acid oxidation resulted in lower voluntary exercise, reduced energy expenditure, and altered adiposity. These data indicate that mildronate-mediated carnitine deficiency in mice provides a means to model and study systemic carnitine deficiency *in vivo*.

## Acknowledgements

We acknowledge the technical support of Trae Pittman, Bobby Fae, Taylor Ross, and Lindsey Pack in the execution of animal studies and sample analyses. This study was supported by NIGMS R35GM142744 and P20GM109096. Support was also provided by the Arkansas Biosciences Institute, the USDA-ARS (USDA ARS 6026-10700-001-000D), and the Barton fund from the University of Arkansas for Medical Sciences. We acknowledge the IDeA National Resource for Quantitative Proteomics for conducting the proteomic work as described in this study (R24GM137786).

## References

1. Gulewitsch W, Krimberg R. Physiol Chem. 1905;45:326–30.

2. Tomita M, Sendju Y. Über die Oxyaminverbindungen welche die Biuret Reaktionen zeigen. III. Spaltung der γ-amino-β-oxybuttersäure in die optisch-aktiven Komponente. Physiol Chem. 1927;169:263–77.

3. Brass EP. Pharmacokinetic considerations for the therapeutic use of carnitine in hemodialysis patients. Clin Ther 1995;17:176–85.

4. Stephens F, Constantin-Teodosiu D, Greenhaff P. New insights concerning the role of carnitine in the regulation of fuel metabolism in skeletal muscle. J Physiol. 2007;581:431–44.

5. Rebouche CJ, Engel AG. Tissue distribution of carnitine biosynthetic enzymes in man. Biochim Biophys Acta. 1980a;5:22–9.

6. Friedman S, Fraenkel G. Reversible enzymatic acetylation of carnitine. Arch Biochem Biophys 1955;59:491–501.

7. Fritz I. The effect of muscle extracts on the oxidation of palmitic acid by liver slices and homogenates. Acta Physiol Scand 1955;12:367–85.

8. Constantin-Teodosiu D, Carlin J, Cederblad G, Harris R, Hultman E. Acetyl group accumulation and pyruvate dehydrogenase activity in human muscle during incremental exercise. Acta Physiol Scand. 1991a;143:367–72.

9. Constantin-Teodosiu D, Cederblad G, Hultman E. PDC activity and acetyl group accumulation in skeletal muscle during prolonged exercise. J Appl Physiol. 1992;73:2403–7.

10. Merritt Jn, Norris M, Kanungo S. Fatty acid oxidation disorders. Ann Transl Med. 2018;6:473.

11. Kerner J, Hoppel C. Genetic disorders of carnitine metabolism and their nutritional management. Annu Rev Nutr 1998;18:179–206.

12. Longo N, Amat di San Filippo C, Pasquali M. Disorders of carnitine transport and the carnitine cycle. Am J Med Genet C Semin Med Genet. 2006;15;142C(2):77–85.

13. Engel A, Angelini C. Carnitine deficiency of human skeletal muscle with associated lipid storage myopathy: a new syndrome. Science. 1973;179:899–902.

14. Treem W, Stanley C, Finegold D, Hale D, Coates P. Primary carnitine deficiency due to a failure of carnitine transport in kidney, muscle, and fibroblasts. N Engl J Med. 1988;319:1331–6.

15. Rebouche CJ, Engel AG. Kinetic compartmental analysis of carnitine metabolism in the human carnitine deficiency syndromes. Evidence for alterations in tissue carnitine transport. J Clin Invest. 1984;73:857–67.

16. Nezu J, Tamai I, Oku A, Ohashi R, Yabuuchi H, Hashimoto N, et al. Primary systemic carnitine deficiency is caused by mutations in a gene encoding sodium ion-dependent carnitine transporter. Nat Genet. 1999;21:91–4.

17. Lennon D, Shrago E, Madden M, Nagle F, Hanson P, Zimmerman S. Carnitine status, plasma lipid profiles, and exercise capacity of dialysis patients: effects of a submaximal exercise program. Metabolism. 1986;35:728–35.

18. Siami G, Clinton M, Mrak R, Griffis J, Stone W. Evaluation of the effect of intravenous L-carnitine therapy on function, structure and fatty acid metabolism of skeletal muscle in patients receiving chronic hemodialysis. Nephron. 1991;57:306–13.

19. Brass EP, Adler S, Sietsema KE, Hiatt WR, Orlando AM, Amato A. Intravenous L-carnitine increases plasma carnitine, reduces fatigue, and may preserve exercise capacity in hemodialysis patients. Am J Kidney Dis. 2001;37:1018–28.

20. Bonafé L, Berger M, Que Y, Mechanick J. Carnitine deficiency in chronic critical illness. Curr Opin Clin Nutr Metab Care. 2014;17:200–9.

21. Murakami K, Sugimoto T, Woo M, Nishida N, Muro H. Effect of L-carnitine supplementation on acute valproate intoxication. Epilepsia. 1996;37:687–9.

22. DeVivo DC. Effect of L-carnitine treatment for valproate-induced hepatotoxicity. Neurology. 2002;58:507–8.

23. Horiuchi M, Kobayashi K, Yamaguchi S, Shimizu N, Koizumi T, Nikaido H, et al. Primary defect of juvenile visceral steatosis (jvs) mouse with systemic carnitine deficiency is probably in renal carnitine transport system. Biochim Biophys Acta. 1994;1226:25–30.

24. Dambrova M, Liepinsh E, Kalvinsh I. Mildronate: Cardioprotective Action through Carnitine-Lowering Effect Trends in Cardiovascular Medicine. 2002;12:275–9.

25. Spaniol M, Brooks H, Auer L, Zimmermann A, Solioz M, Stieger B, et al. Development and characterization of an animal model of carnitine deficiency. Eur J Biochem. 2001;268:1876–87.

26. Sadler DG LR, Treas L, Sikes JD, Porter C. Protonophore treatment augments energy expenditure in mice housed at thermoneutrality. Protonophore treatment augments energy expenditure in mice housed at thermoneutrality. Frontiers in Physiology. 2024;In press.

27. Sadler DG TL, Sikes JD, Porter C. A modest change in housing temperature alters whole body energy expenditure and adipocyte thermogenic capacity in mice. Am J Physiol Endocrinol Metab. 2022;323(6):E517–E28.

28. Ge SX, Jung D, R Y. ShinyGO: a graphical gene-set enrichment tool for animals and plants. Bioinformatics. 2020;36(8):2628–9.

29. Conway JR, Lex A, N G. UpSetR: an R package for the visualization of intersecting sets and their properties. Bioinformatics. 2017;33(18):2938–40.

30. Porter C, Constantin-Teodosiu D, Constantin D, Leighton B, Poucher S, Greenhaff P. Muscle carnitine availability plays a central role in regulating fuel metabolism in the rodent. J Physiol. 2017;595:5765–80.

31. Simkhovich B, Shutenko Z, Meirena D, Khagi K, Mezapuķe R, Molodchina T, et al. 3-(2,2,2-Trimethylhydrazinium)propionate (THP)--a novel gamma-butyrobetaine hydroxylase inhibitor with cardioprotective properties. Biochem Pharmacol 1988;15:195–202.

32. Kuwajima M, Harashima H, Hayashi M, Saori I, Masako S, Kang-Mo L, et al. Pharmacokinetic analysis of the cardioprotective effect of 3-(2,2,2- trimethylhydrazinium) propionate in mice: Inhibition of carnitine transport in kidney. Journal of Pharmacology and Experimental Therapeutics 1999;289:93–102.

33. Georges B, Le Borgnea F, Gallanda, Isoira M, Ecossea D, Grand-Jeana F, et al. Carnitine transport into muscular cells. inhibition of transport and cell growth by mildronate Biochemical Pharmacology. 2000;59:1357–63.

34. Grigat S, Fork C, Bach M, Golz S, Geerts A, Schömig E, et al. The carnitine transporter SLC22A5 is not a general drug transporter, but it efficiently translocates mildronate. Drug Metab Dispos. 2009;37:330–7.

35. Liepinsh E, Kuka J, Svalbe B, Vilskersts R, Skapare E, Cirule H, et al. Effects of Long-Term Mildronate Treatment on Cardiac and Liver Functions in Rats. Basic Clin Pharmacol Toxicol. 2009a;105:387–94.

36. Liepinsh E, Vilskersts R, Zvejniece L, Svalbe B, Skapare E, Kuka J, et al. Protective effects of mildronate in an experimental model of type 2 diabetes in Goto-Kakizaki rats. Br J Pharmacol 2009b;157:1549–56.

37. Tsoko M, Beauseigneura F, Grestia J, Niota I, Demarquoya J, Boichota J, et al. Enhancement of activities relative to fatty acid oxidation in the liver of rats depleted of -carnitine by -carnitine and a γ-butyrobetaine hydroxylase inhibitor 1995;49:1403–10

38. Roberts P, Bouitbir J, Bonifacio A, Singh F, Kaufmann P, Urwyler A, et al. Contractile function and energy metabolism of skeletal muscle in rats with secondary carnitine deficiency. Am J Physiol Endocrinol Metab. 2015;309:265–74.

39. Bouitbir J, Haegler P, Singh F, Joerin L, Felser A, Duthaler U, et al. Impaired Exercise Performance and Skeletal Muscle Mitochondrial Function in Rats with Secondary Carnitine Deficiency. Front Physiol. 2016;7(345).

40. Whitener D, Whitener L, Robertson K, Baxter C, Pierce A. Pulmonary function measurements in patients with thermal injury and smoke inhalation. Am Rev Respir Dis. 1980;122:731–9.

41. Stephens F, Wall B, Marimuthu K, Shannon C, Constantin-Teodosiu D, Macdonald I, et al. Skeletal muscle carnitine loading increases energy expenditure, modulates fuel metabolism gene networks and prevents body fat accumulation in humans. J Physiol. 2013;591(18):4655–66.

42. Stephens FB, Constantin-Teodosiu D, Laithwaite D, Simpson EJ, Greenhaff PL. An acute increase in skeletal muscle carnitine content alters fuel metabolism in resting human skeletal muscle. J Clin Endocrinol Metab. 2006;91:5013–8.

43. Wall B, Stephens F, Constantin-Teodosiu D, Marimuthu K, Macdonald I, Greenhaff P. Chronic oral ingestion of L-carnitine and carbohydrate increases muscle carnitine content and alters muscle fuel metabolism during exercise in humans: the dual role of muscle carnitine in exercise metabolism. J Physiol. 2011.

44. Pereyra AS, McLaughlin KL, Buddo KA, JM. E. Medium-chain fatty acid oxidation is independent of l-carnitine in liver and kidney but not in heart and skeletal muscle. Am J Physiol Gastrointest Liver Physiol. 2023;325(4):G287–G94.

45. Sidossis L, Stuart C, Shulman G, Lopaschuk G, Wolfe R. Glucose plus insulin regulate fat oxidation by controlling the rate of fatty acid entry into the mitochondria. J Clin Invest. 1996;98:2244–50.

46. Rasmussen B, Holmbäck U, Volpi E, Morio-Liondore B, Paddon-Jones D, Wolfe R. Malonyl coenzyme A and the regulation of functional carnitine palmitoyltransferase-1 activity and fat oxidation in human skeletal muscle. J Clin Invest. 2002;110:1687–93.

47. Hiatt W, Wolfel E, Regensteiner J, Brass E. Skeletal muscle carnitine metabolism in patients with unilateral peripheral arterial disease. J Appl Physiol. 1992; 73:346–53.

48. Tein I. Carnitine transport: pathophysiology and metabolism of known molecular defects. J Inherit Metab Dis. 2003;26:147–69.

